# On the preservation of vessel bifurcations during flow-mediated angiogenic remodelling

**DOI:** 10.1101/2020.02.07.938522

**Authors:** Lowell T. Edgar, Claudio A. Franco, Holger Gerhardt, Miguel O. Bernabeu

## Abstract

During developmental angiogenesis, endothelial cells respond to shear stress by migrating and remodelling the initially hyperbranched plexus, removing certain vessels whilst maintaining others. The key regulator of vessel preservation is cell decision behaviour at bifurcations. At flow-convergent bifurcations where migration paths diverge, cells must finely tune migration along both possible paths if the bifurcation is to persist. Experiments have demonstrated that disrupting the cells’ ability to sense shear or junction forces transmitted between cells impacts the preservation of bifurcations during the remodelling process. However, how these migratory cues integrate during cell decision making remains poorly understood. Therefore, we present the first agent-based model of endothelial cell flow-mediated migration suitable for interrogating the mechanisms behind bifurcation stability. The model simulates flow in a bifurcated vessel network composed of agents representing endothelial cells arranged into a lumen which migrate against flow. Upon approaching a bifurcation where more than one migration path exists, agents refer to a stochastic bifurcation rule which models the decision cells make as a combination of flow-based and collective-based migratory cues. With this rule, cells favour branches with relatively larger shear stress or cell number. We found that cells must integrate both cues nearly equally to maximise bifurcation stability. In simulations with stable bifurcations, we found competitive oscillations between flow and collective cues, and simulations that lost the bifurcation were unable to maintain these oscillations. The competition between these two cues is haemodynamic in origin, and demonstrates that a natural defence against bifurcation loss during remodelling exists: as vessel lumens narrow due to cell efflux, resistance to flow and shear stress increases, attracting new cells to enter and rescue the vessel from regression. Our work provides theoretical insight into the role of junction force transmission has in stabilising vasculature during remodelling and as an emergent mechanism to avoid functional shunting.

**Author Summary:** When new blood vessels are created, the endothelial cells that make up these vessels migrate and rearrange in response to blood flow to remodel and optimise the vessel network. An essential part of this process is maintaining the branched structure of the network; however, it is unclear what cues cells consider at regions were vessels branch (i.e., bifurcations). In this research, we present a computer model of cell migration to interrogate the process of preserving bifurcations during remodelling. In this model, cells at bifurcations are influenced by both flow and force transmitted from neighbouring cells. We found that both cues (flow-based and collective-based) must be considered equally in order to preserve branching in the vessel network. In simulations with stable bifurcations, we demonstrated that these cues oscillate: a strong signal in one was accompanied by a weak signal in the other. Furthermore, we found that these cues naturally compete with each other due to the coupling between blood flow and the size of the blood vessels, i.e. larger vessels with more cells produce less flow signals and vice versa. Our research provides insight into how forces transmitted between neighbouring cells stabilises and preserves branching during remodelling, as well as implicates the disruption of this force transmission as a potential mechanism when remodelling goes wrong as in the case of vascular malformation.

## Introduction

Angiogenesis occurs as two distinct phases: an early phase in which sprouting neovessels assemble to form the initial immature vascular plexus, and a late phase which remodels the plexus into its final functional form [1,2]. Blood flow is typically shunted away from sprouting endothelial cells (ECs), which exhibit a dynamic exploratory phenotype as they invade the avascular space. However, ECs exhibit a fundamentally different phenotype upon receiving blood flow: in response to experiencing the physical force of flow (i.e., wall shear stress along the luminal surface, WSS), ECs will re-align their polarity and migrate against the direction of flow [3–8]. This phenomenon, referred to as EC flow-migration coupling [8], has been shown to be the primary driver of vascular remodelling during developmental angiogenesis.

There are two principal outcomes of flow-mediated remodelling: the pruning of superfluous connections that arise during the sprouting phase, and the establishment of vascular hierarchy via diameter control. Under the flow-migration hypothesis, ECs move from low-flow vessels as they are attracted to high-flow vessels along a path opposite to flow. This attraction results in pruning of inefficient vessels while reinforcing the established preferred flow paths, including precursors to arteries and veins. This hypothesis presents a paradigm shift through which we can now view quiescent vascular structures as nonlinear dynamic systems of migrating cells which have reached stability at a critical point and shifts between healthy vascular tissue and diseased now as transitions between stable fixed points which may be reversible. However, the fundamental question as to how vessel networks determine the appropriate structure and hierarchy needed to establish optimal tissue perfusion during flow-regulated migration and remodelling remains largely unanswered.

Branching within the vasculature is a key component to effective transport, and it appears that an important result of flow-mediated remodelling is to remove some complexity from the initial plexus in order to improve efficiency while maintaining enough complexity within the network to ensure large surface area and short diffusion distance to the surrounding tissue. However, under the basic flow-migration hypothesis, in which ECs simply move against flow from areas of low-flow to high, a discrepancy arises at bifurcations: what do ECs do at flow-convergent bifurcations in which two paths to migrate against flow exist? Do ECs simply choose the high-flow branch, thereby reinforcing these vessels at the expense of their low-flow counterparts? Or does some mechanism exist that allows bifurcations with a flow difference between the two branches to persevere during remodelling? The flow-migration hypothesis is relatively new in its development and lacks a description of EC behaviour at vessel bifurcations as well as consideration as to how certain bifurcations are preserved and not others.

Very little is known about the mechanisms regulating EC flow-migration coupling; however, some signalling pathways have been revealed to be involved. In particular, the noncanonical Wnt signalling pathway plays a profound role. Franco et al. demonstrated that knocking down Wnt5a in mutant mice (hence referred to as Wnt KD) rendered ECs in the developing retina more sensitive to WSS, increasing their axial polarisation against flow and migration levels [6]. Additionally, they found reduced amounts of bifurcations and increased regression events within these mice, indicating that increased levels of flow-mediated remodelling reduced the branching complexity of the emerging vasculature. Noncanonical Wnt signalling has also been implicated in sprouting ECs as well in a different role. Carvalho et al. demonstrated the role of Wnt5a in collective cell behaviour at the sprouting front, where Wnt5a works to reinforce leader-follower collective polarity by strengthening and stabilising adherens junctions, facilitating force transmission between ECs [9].

Putting these findings together demonstrates that interfering with noncanonical Wnt5a signalling reduces junction force transmission between ECs, which facilitates better flow-based polarisation and migration, increases vascular remodelling, and results in a network that is less branched. The fact that reducing junction force transmission facilitates flow-based remodelling rather than inhibits may seem counterintuitive, as many collective migration processes depend on junction force transmission [10,11]. Rather, it would seem that flow-migration coupling during remodelling is a different type of process all together, and the “collectiveness” of the ECs interferes with the process in order to keep remodelling in check. This implies that flow-migration coupling is more of an “individualistic” process in which isolated ECs responding to a global signal (i.e., WSS due to blood flow), rather than a process regulated via cell-cell communication.

We hypothesise that EC migration during angiogenic remodelling results from integrating both flow-directed migration (guided by differences in luminal shear stress) and collective migration (guided by cell-to-cell junctional force transmission). Furthermore, we propose that these two migratory cues interact (either cooperatively or competitively) to determine bifurcation stability and promote or avoid vessel regression during remodelling. In this work we present an agent-based model (ABM) of EC flow-migration coupling to demonstrate that competition between flow-based and collective-based cues can stabilise vessel bifurcations during remodelling. ABMs have proven to be useful tools in the study of sprouting angiogenesis due to their naturally discrete representation of cells and their ability to capture emergent behaviour [12–20]; however, there are currently no models of the coupling between flow and migration during remodelling to date. In ABMs, we can prescribe various “rules” of ECs behaviour at bifurcations and observe the outcome as an emergent property, allowing us to characterise the impact of different mechanisms when it comes to achieving bifurcation stability. We found that bifurcation stability cannot be achieved when ECs follow flow or collective migration cues alone. The best outcome for bifurcation stability was when we included near equal contributions of shear stress and junction force transmission cues when determining EC migration at bifurcations. Additionally, we found that this stability arises due to the competitive interplay of the two mechanisms, which oscillate back in forth to keep the bifurcation stable. The competitive nature of these two cues is of hydrodynamic origin, arising from the inverse relationship between lumen diameter and vessel resistance to flow and can be characterised by a single parameter. We postulate that published observations on the molecular regulation of EC flow-migration coupling can be re-envisioned under this new lens, offering a powerful rationale for developing a mechanistic understanding of vascular remodelling dynamics at the level of the whole vascular plexus.

## Results

### Agent-based model of EC migration coupled to flow

In this study we present an agent-based model of EC migration coupled to blood flow within an idealised bifurcated vessel network (hence referred to as the A branch, see the Methods Section for a complete description). This network consists of a feeding vessel and a draining vessel, connected by a proximal branch and a distal branch (Fig. 1 A). Each vessel was discretised into segments and each segment was seeded with an initial number of “agents” representing ECs (Fig. 1 B). Flow was driven by the difference in pressure prescribed at the inlet and outlet in order to recapitulate previously reported WSS values [21], and calculated via the Hagen–Poiseuille equation while treating blood as a Newtonian Fluid with constant viscosity. The flow conductance (i.e., the inverse of the flow resistance) was calculated using a three-dimensional (3D) approximation of the lumen diameter by wrapping the number of cells in each vessel segment into the circumference of a circle (Fig 1 C). Each simulation was run for a prescribed number of steps in time, and during each step ECs moved against the direction of flow to the neighbouring upstream segment. Periodic boundary conditions were prescribed to handle the case of ECs at the inlet, in which case they migrated to the outlet (thus preserving the total number of cells during each simulation). After each time step, vessel diameter was updated depending on the new number of ECs in each segment and flow re-calculated, thus directly linking flow with migration.

**Fig 1.**
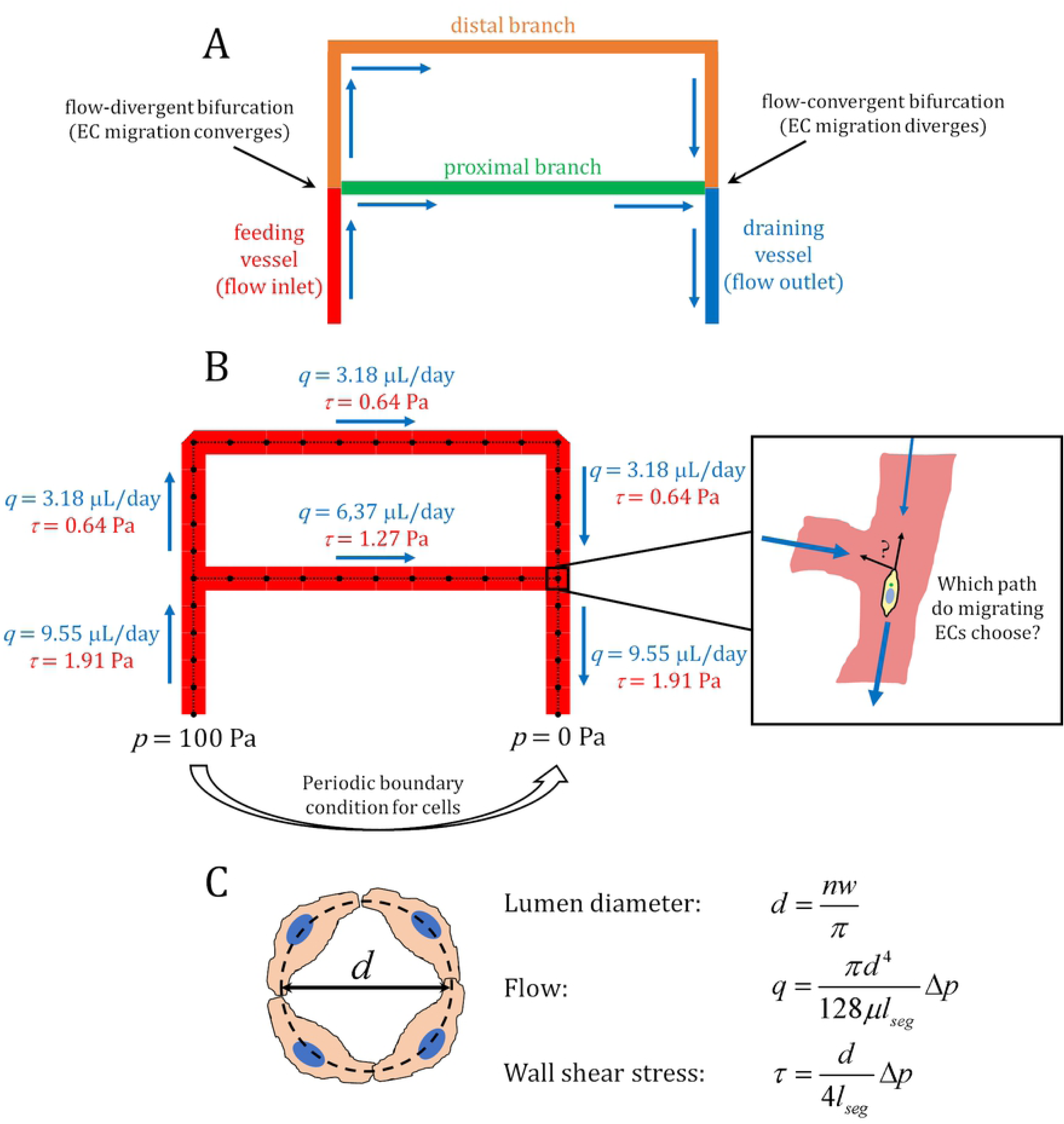
Flow-migration coupling within the A branch model. An idealised model of a vessel bifurcation with shear stress differences present. (A) The network consists of a feeding vessel connected to a draining vessel by a short proximal path and a longer distal path. Blue arrows indicate the direction of flow throughout the network. Flow at the bifurcation near the inlet diverges, while flow near the outlet converges. The difference in path lengths results in different levels of flow/shear stress within each branch. (B) The network was discretised and seeded with an initial number of ECs (agents). Pressure boundary conditions (black) and values of flow (blue) and shear stress (red) are given in the initial configuration of the network. Periodic boundary conditions were prescribed at the inlet and outlet in order to keep the total number of cells within the simulation constant. In this model we are concerned with EC behaviour at flow-convergent bifurcations where two options to migrate against the flow exist: which path to the migrating ECs choose, and what determines this choice? (C) Vessel lumens are approximated in 3D by wrapping the number of cells in the vessel, *n*, each with width *w*, into the circumference of a circle. Flow and shear stress are then calculated using the Hagan-Poiseuille equation.

The network contains two different types of bifurcations: a flow-divergent bifurcation at the feeding vessel where the flow splits between the proximal and distal branch, and a flow-convergent bifurcation at the draining vessel where the flow from the branches combine before draining at the outlet. Additionally, the difference in lengths between the proximal and distal paths results in an initial shear stress difference at the bifurcations. This network configuration provides a simple and systematic setting to investigate the consequences of EC behaviour at bifurcations during flow-mediated migration and remodelling where shear stress differences are present. Bifurcations where flow diverges are areas where EC migration paths converge, and flow-convergent bifurcations where ECs paths diverge. Migrating ECs approaching the flow-divergent bifurcation from either branch have only one option to continue migrating against the flow (i.e., to enter the feeding vessel towards the inlet). However, at the flow-convergent branch, two paths against the flow exist: either into the high-flow proximal branch, or low-flow distal branch. The essence of our current research effort is to investigate how ECs choose which path to follow at these bifurcations, and what the consequences of such choices on vascular remodelling may be.

### Wall shear stress alone cannot stabilise bifurcations

Our initial investigation involved implementing a series of rules for behaviour at the bifurcation (or bifurcations rules, BRs) and observed the emergent outcome in the resulting simulations. In the first bifurcation rule (BR 1), we programmed that upon reaching the flow-convergent bifurcation, the EC would choose the branch with larger shear stress (Fig 2 A, S1 File). This simulation always resulted in the loss of the low-flow distal branch, as cells would always turn into the high-flow branch, meaning cells migrating out of the distal branch were not replenished with new cells. This resulted in a loss of the bifurcation within the network as the cells formed into a single path from inlet to outlet along the proximal branch. Turning into the high-flow branch as in BR 1 means that cells would have to make a sharp change in their migration direction. Therefore, in BR 2 we interrogated if cells would prefer the path that required the smallest change in direction (Fig 2 B, S2 File). This simulation always results in the loss of the proximal branch and reinforcement of the distal branch, as this path requires no direction change from ECs at the bifurcation.

**Fig 2.**
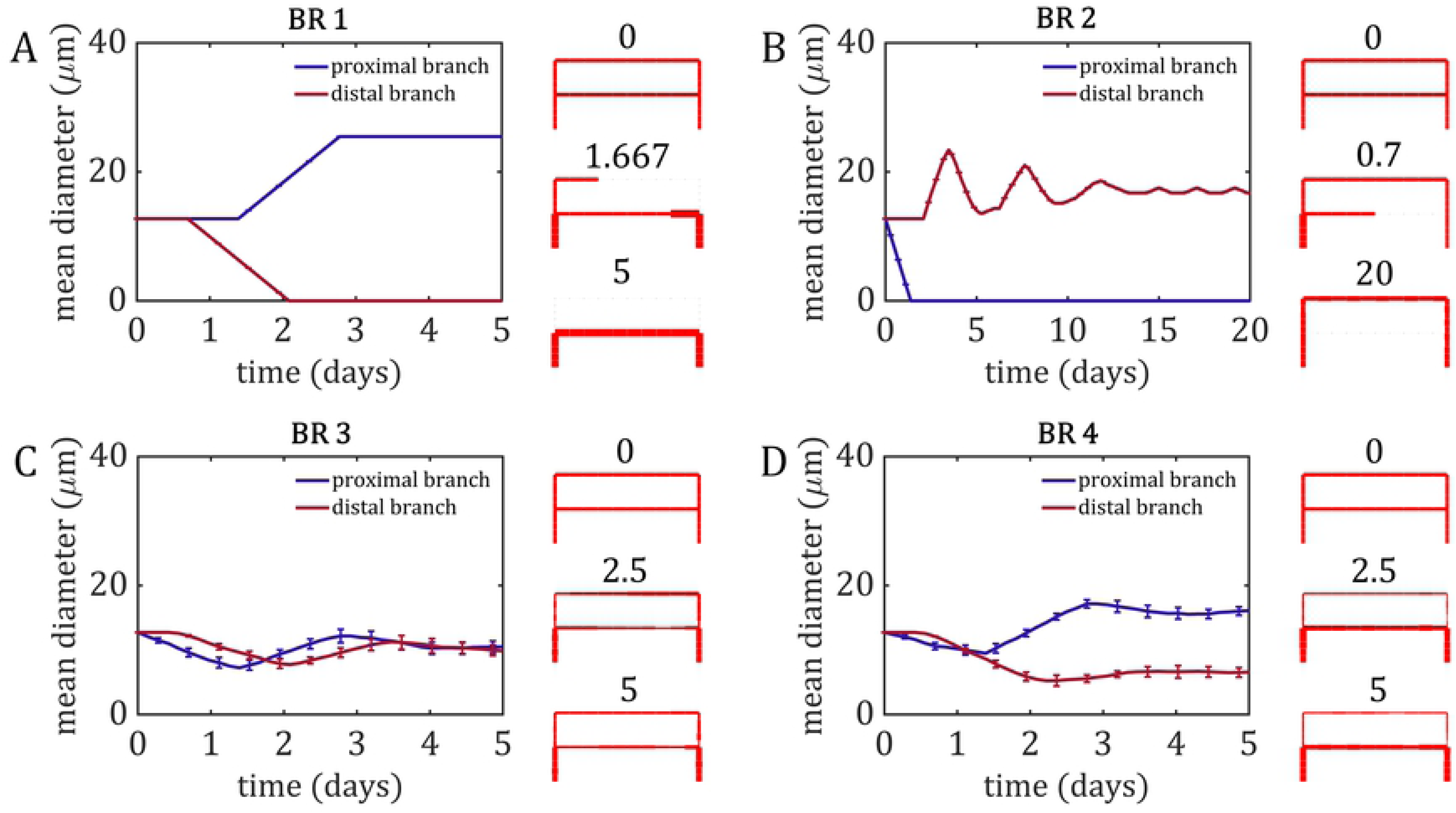
Simple bifurcation rules (BRs 1 through 4). The mean diameter was calculated over all segments composing the proximal branch (blue) and the distal branch (red). Snapshots of the network at various time points are presented on the far right of every panel. (A) Simulations using BR 1 always resulted in the regression of the distal branch and bifurcation loss, as all cells chose to enter the high-flow proximal branch. (B) In BR 2, cells always chose the path that requires the smallest change in migration direction, resulting in the loss of the proximal branch and the bifurcation. This simulation experiences numerous oscillations in diameter as the distal path is twice the length of the proximal path, and it takes several trips around before the smoothing algorithm settles down the diameter fluctuations. (C) Mean diameter and standard deviation for the 10 runs using BR3.. Using a random number generator to determine which branch each cell entered with equal probability resulted in stabilisation of both branches and no discernible difference in diameter between the two branches. (D) Mean diameter and results of each run using BR 4. Using unequal probability between the two branches, while favouring the high-flow branch, resulted in a form of diameter control, as the high-flow proximal branch stabilised at a larger diameter than the low-flow distal branch.

We then asked the question of how the flow-convergent bifurcation may remain in such a network configuration. It would seem that for the bifurcation to remain stable, some of the cells would have to choose the high-flow option, while others choose the path requiring the least change in direction/polarisation. Therefore, we implemented a new mechanism in which each branch was assigned equal probability of drawing in new cells (*P*_1_ = *P*_2_ = 0.5, BR 3). In these simulations, both the proximal and distal branch remained stable, preserving the bifurcation, and stabilised at a similar diameter despite the difference in flow between them (Fig 2 C, S3 File). Finally, it is known that diameter control is an important emergent outcome of vascular remodelling as high-flow vessels become stable at larger diameters than vessels with lower flow, establishing vascular hierarchy. Therefore, we modified the previous bifurcation rule so that the high-flow segment had a higher fixed probability of drawing in new ECs than the low-flow segment (*P*_1_ = 0.7; *P*_2_ = 0.3, BR 4). In these simulations, both vessels remained stable and the bifurcation was preserved, but the high-flow proximal branch stabilised at a larger diameter than the low-flow distal branch (Fig 2 D, S4 File).

Using BRs 3 and 4, we were able to preserve the branched structure of the vascular network as well as render a form of diameter control during flow-mediated EC migration. However, our simple bifurcation rules were based solely on randomness and arbitrarily chosen probabilities and lack a mechanism as to how and why bifurcations remain stable. Therefore, we took steps to create a bifurcation rule which contained more physiologically relevant mechanisms based on what we know about flow-mediated vascular remodelling *in vivo*. As mentioned previously, experiments involving noncanonical Wnt5a signalling suggest competitive interplay between flow-based polarisation/migration and collective cell junction communication. Based on these observations, we designed a mechanistic bifurcation rule (BR 5) through which the probability of each branch “attracting” incoming cells is given by a weighted average of two probability components: one due to shear stress (flow-migration term), and one due to cell number/junction force transmission (collective cell behaviour term) (see Methods, Eq 12-14). We define the probability of an EC choosing a branch due to shear stress as that branch’s contribution to the shear stress ratio (as ECs are attracted towards regions of higher shear). Likewise, probability due to junction forces is defined as the branch’s contribution of the cell number ratio (as the resulting net force direction will favour the branch with larger number of cells). Use of ratios to define probability ensures that our probability definitions always rest between the required range of 0 and 1. The strength of influence each component has on EC decisions is governed by the parameter *α*: shear stress probability scales relative to *α*, and cell number probability relative to 1-*α*. Unlike in BR1 and BR2, ECs will choose the preferred direction of migration only probabilistically (i.e., it is still possible for an EC to choose the non-preferential direction of migration) since we want to investigate stochastic effects arising due to the relatively low number of cells in these vessel networks.

We tested this new bifurcation rule by running the model 1000 times, each with a different random seed, for different values of *α* (using the same 1000 random seed numbers for each value of *α*). Setting *α* to 0.0 means that only collective cell behaviour is considered at the bifurcation; 91.9% of simulations at this value of *α* experienced regression and bifurcation loss with no obvious preference for the proximal or distal branch (Fig 3 A, S5 File). A value of *α* = 1.0 means that only shear stress is considered at the bifurcation; these simulations were even more unstable, with 100% resulting in bifurcation loss shortly after 2 days of migration (Fig 3 B, S6 File). Additionally, in these simulations the low-flow distal branch was 2.7× more likely to regress than the high-flow proximal branch. Simulations at this value of *α* were prone to lose the bifurcation much sooner than their counterparts at *α* = 0.0, which didn’t reach over 90% loss until 5 days of migration. It should be noted that in this bifurcation rule, branch probability continuously adapts to changes in flow and cell number (rather than being held constant as seen in BRs 3 and 4). This results in rapid adaptation of the vasculature to the various stochastic outcomes which can push the model to stable solutions that may seem unintuitive *a priori*. For example, in nearly 25% of cases with *α* = 1.0 resulted in loss of the proximal branch in favour of the longer distal path, despite the proximal path initially experiencing larger shear stress. These results demonstrate the complex nature of the system, which can quickly and dramatically push towards different stable outcomes given accumulation of small stochastic effects.

**Fig 3.**
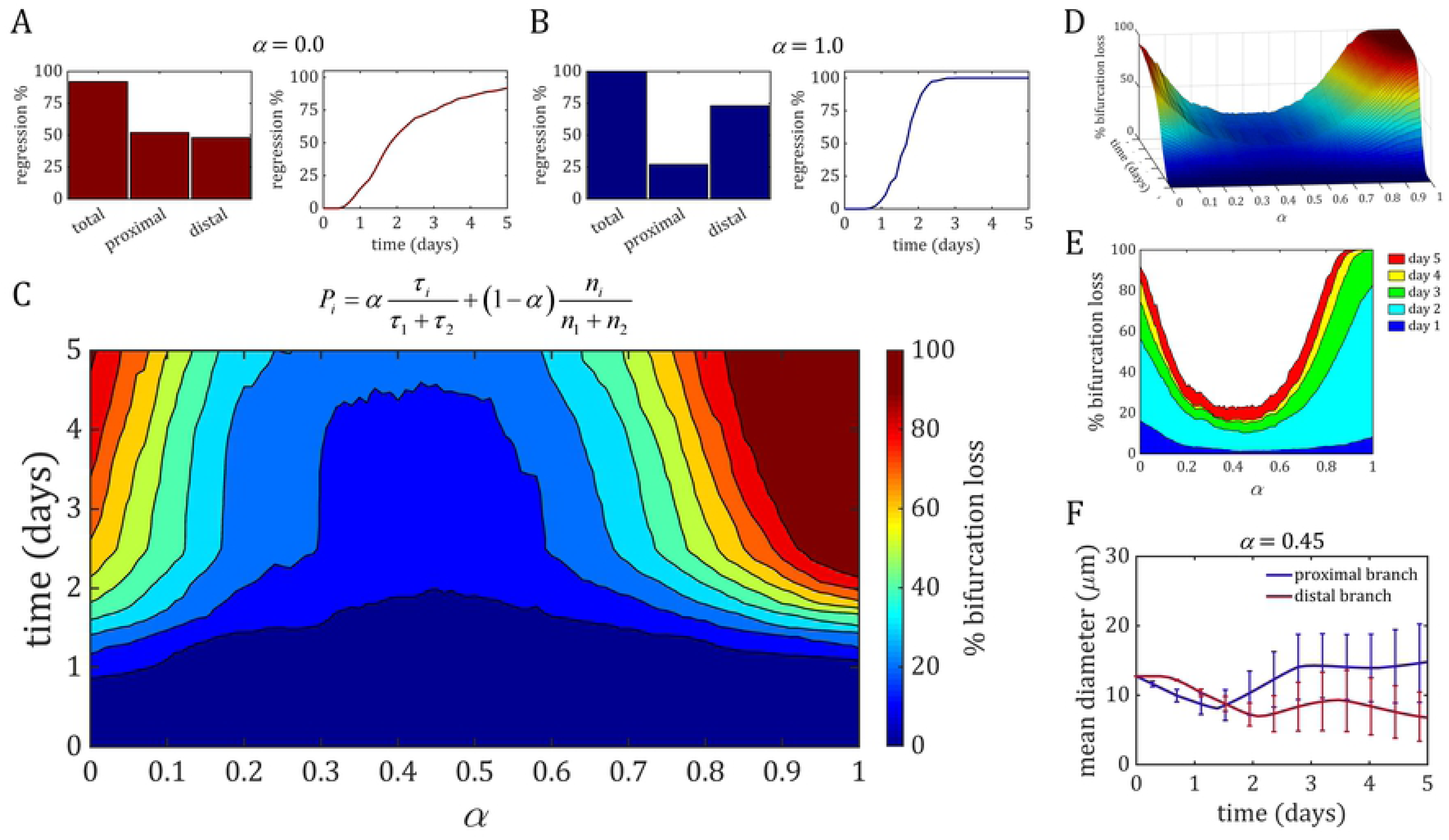
Stability analysis of the weight parameter *α* in the mechanistic bifurcation rule. The weight parameter *α* is used to scale the respective influence of the shear stress ratio and cell number ratio when calculating probability of a cell to enter each branch. We discretised *α* over its range. (A) On the left, the percentage of total regression events in all simulations and the percentage of those regression events that involved either the proximal or distal branch. On the right, the percentage of regression events over time. Setting *α* = 0.0 means only cell number was used to determine the probability of each branch; 91.9% of these simulations experienced regression and bifurcation loss, with 52% involving proximal regression vs. 48% distal (B) Setting *α* = 1.0 and using only shear stress to determine branch probability resulted in 100% of simulations losing the bifurcation, with 27% involving proximal regression vs. 73% distal. Additionally, these simulations were prone to lose the bifurcation much earlier, with near 100% regression reached after just 2 days of migration. (C) Contour plot of bifurcation loss over the whole range of *α*, which demonstrates a global minimum of stability for values of *α* ranging from 0.3 to 0.6. (D) The surface of bifurcation loss vs. *α* over time resembles and asymmetric saddle with similar rates of increase in bifurcation loss in both directions. (E) Bifurcation loss grouped within each day of migration. The majority of bifurcations were lost during day 2, while minimal loss occurred after that. (F) Mean diameter of the proximal (blue) and distal branch (red) with standard deviation for simulations within the stable region (*α* = 0.45). The high-flow proximal branch, on average, stabilised at a larger diameter than the low-flow distal branch. Note that both the proximal and distal branch experience some initial transients in cell number/diameter which eventually stabilise once cells have traversed each path completely. As a result, transients take longer to settle in the distal branch due to the longer path length.

The question now is: what is an optimal value of *α* in order to maximise bifurcation stability? To determine the performance of *α*, we swept through its range (from 0.0 to 1.0 at increments of 0.01) and ran the 1000 seeds of the random number generator. We determined if and at what point in time the bifurcation was lost (declared when cell number in one of the two branches at the flow-convergent bifurcation dropped to zero) and summed this value for all runs of each value of *α*. Dividing this number by the number of runs (*M* = 1000) provides us the percentage of simulations in which the bifurcation was lost over time (Fig 3 C). In general, simulations which favoured shear stress when determining bifurcation behaviour (*α* > 0.5) where much more prone to bifurcation loss than simulations favouring collective cell behaviour (*α* < 0.5). The most stable values of *α* were between 0.3-0.6, with a peak of stability centred around *α* = 0.45. The surface formed by bifurcation loss vs. *α* over time resembles an asymmetric saddle that flattens out in the ranges of *α* between 0.3 and 0.6 (Fig. 3 D). From this surface it is apparent that the rate of bifurcation loss as *α* approaches either 0.0 or 1.0 is similar; however, simulations with values of *α* > 0.5 become unstable earlier and reach the plateau sooner than values of *α* < 0.5. When observing the cross-section of the surface at different days, it seems that most of the bifurcation loss occurs over day 2, with minimal loss at later days (Fig 3 E). Interestingly, values of *α* = 0.0 were slightly more unstable over day 1 than values of *α* = 1.0; however, these simulations became much more unstable at later days compared to their counterparts. Finally, when looking at mean diameter of both the proximal and distal branch at what appears to be peak stability at *α* = 0.45, only 22.9% of these simulations experienced bifurcation loss with the distal branch 5× more likely to regress compared to the proximal branch. Mean diameter over time shows that these simulations preferred to stabilise the high-flow proximal branch at a larger diameter than the low-flow distal branch (Fig. 3 F, S7 File).

### Competitive oscillations between flow-based and collective-based mechanisms achieve bifurcation stability

Next, we interrogated simulations at stable values of *α* in order to determine the source of this stability by monitoring the probabilities of the proximal branch (*P*_1_) and the distal branch (*P*_2_). These branch probabilities can be decomposed into a linear combination of the shear stress and cell number components (*P_τ_*_1_, *P_n_*_1_ for the proximal branch and *P_τ_*_2_, *P_n_*_2_ for the distal branch), weighted by the parameter *α* (Fig. 4 A). In simulations at stable values of *α* (e.g., *α* = 0.45), the probability of each branch oscillated slightly over time but centred around a relatively constant probability value (Fig. 4 B). This averaged to a higher value in the high-flow branch (typically between 0.6-0.7) compared to the low-flow branch (between 0.3-0.4), which stabilises this branch at a larger diameter and higher cell number than its low-flow counterpart. When looking at the shear stress and cell number components of these probabilities we find alternating peaks in magnitude between *P_τi_* and *P_ni_* in both branches (Fig. 4 C). Each peak in shear stress probability is accompanied by a trough in cell number probability and vice versa, and every peak in one is both proceeded and followed by a peak in the other. Additionally, every trough in either shear stress or cell number probability in the high-flow branch is accompanied by a peak in the same probability value in the low-flow branch. In simulations that lost the bifurcation, we found temporary oscillations between the two probability components while the bifurcation was stable, but in each of these cases the system was pushed to completely favour one branch over the other (i.e., a branch’s probability equal to 1 while the other equal to 0) resulting in bifurcation loss (S8 Fig).

**Fig 4.**
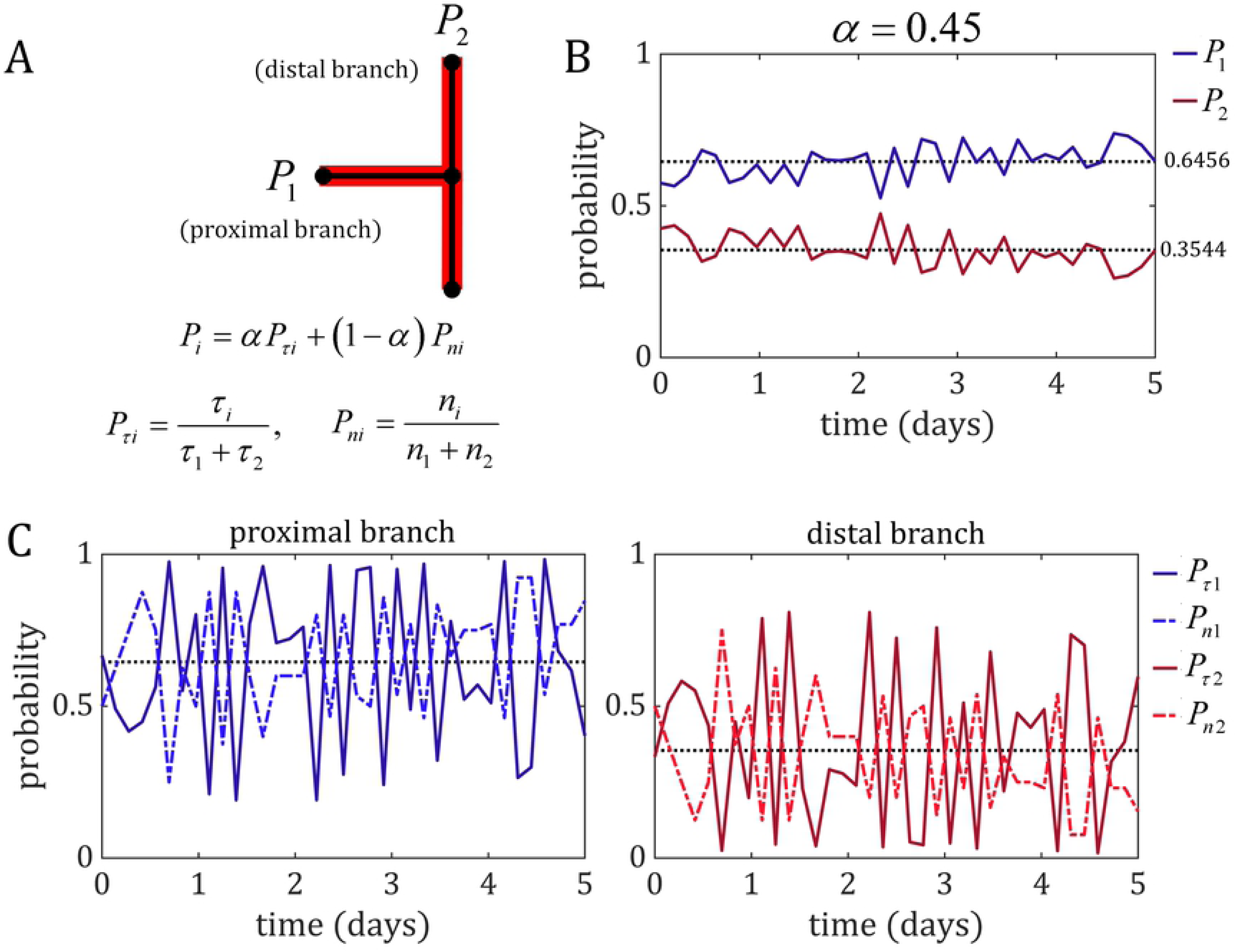
Competitive oscillations between the flow-based mechanism (shear stress) and collective-based mechanism (junction forces) achieves bifurcation stability. (A) The probability of each branch of pulling in new cells at the bifurcation, *P_i_*, is given by the weighted average of the shear stress probability *^P^τi* and the cell number probability *P_ni_*. (B) The branch probability over time for an example stable simulation with a value of *α* = 0.45. The branch probability at the bifurcation oscillated slightly over time, with the probability of the high-flow proximal branch averaging to a higher value (dark blue, mean 0.6456) than the low-flow distal branch (dark red, mean 0.3544). (C) The individual components of these probabilities also oscillate over time in both the proximal branch (left) and the distal branch (right), with alternating peaks in shear stress (dark blue/red, solid line) and cell number probability (light blue/red, dashed-dot line). Additionally, a peak in either component in one branch was accompanied by a trough the that same probability component in the other branch. These data suggest that the shear stress and cell number components interact competitively at stable bifurcations, each compensating for increases in the other to prevent either of the two components from dominating the bifurcation (which would result in a loss of one of the two branches).

The shear stress within each vessel depends on the number of cells in the vessel and the pressure drop across the segment (see Methods, Eq 6). How then is *P_τ_* able to compete with *P_n_* when they both function on the number of cells in the vessel? The pressure drop is tied to the vessel’s resistance to flow: if the vessel narrows, then a larger pressure drop will be required to produce the same amount of flow. However, the resistance is inversely proportional to the number of cells quatrically, *R* ∝ *n*^−4^ (see Methods, Eq 3), and therefore small changes in cell number will quickly manifest as large changes in resistance and the pressure drop required to push incoming flow from upstream segments will similarly increase. The shear stress probability, *P_τ_*, can be simplified to a multiplicative combination of the cell number (∝ *n*^1^) and the pressure drop across the vessel (∝ *n*^−4^) (see Methods, Eq 15). Due to the large difference in the powers of *n*, *P_τ_* is rendered inversely proportional to *n* with a power greater than 1. This means that small fluctuations in cell number lead to large fluctuations in pressure drop, and indeed if we monitor the pressure drop and cell number within the branches at the bifurcation, we find that these two quantities are inversely related with dramatically different magnitudes (Fig. 5).

**Fig 5.**
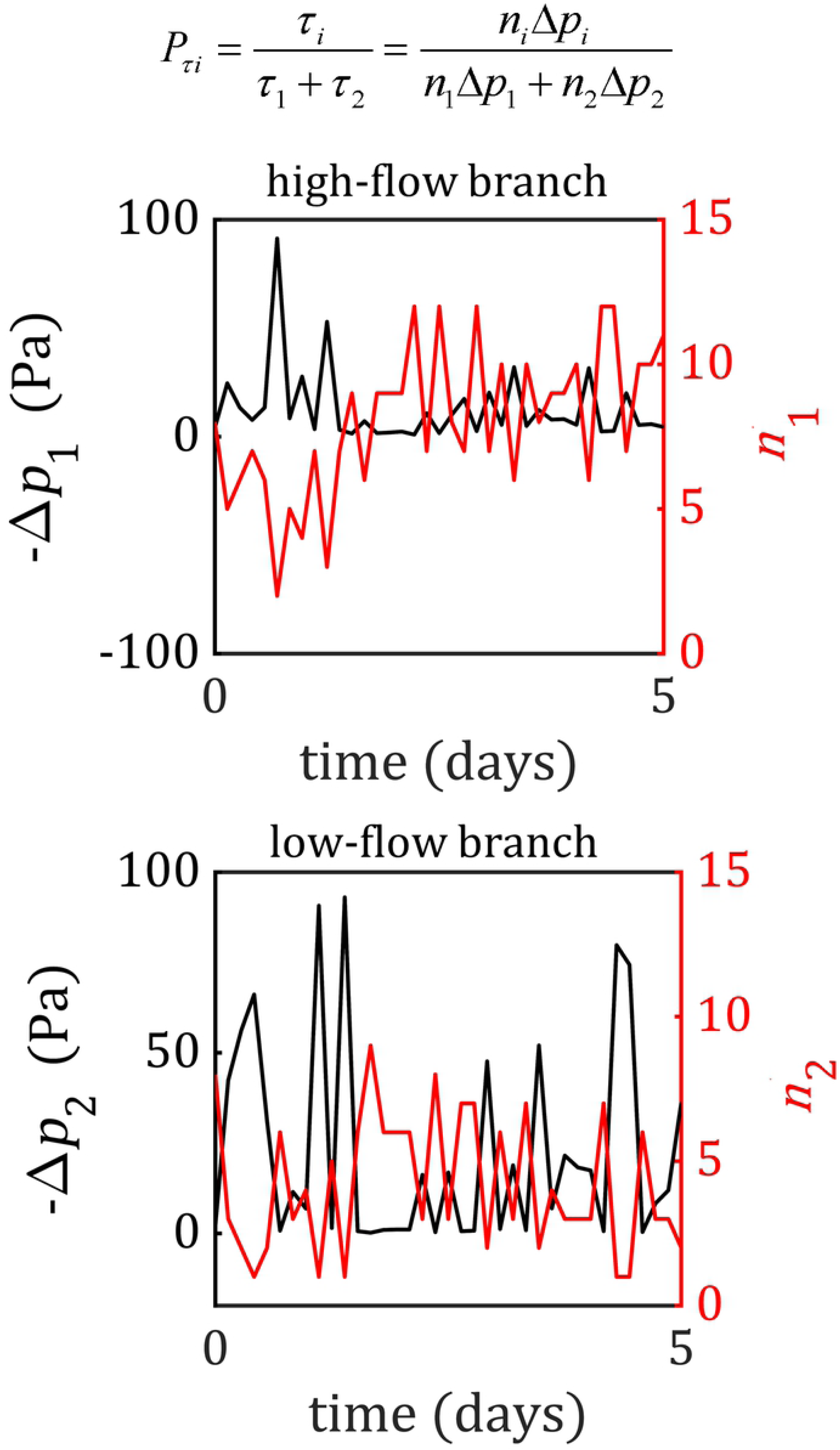
The competition between shear stress and junction force transmission results from the inverse relationship between cell number and flow resistance. The shear stress ratio within each vessel can also be expressed as a multiplicative combination of the cell number and pressure drop over the vessel. If we monitor cell number (red) and the pressure drop (black) in both the high-flow (top) and low-flow (bottom) vessel, we find that these two quantities are inversely correlated. Drops in the cell number are accompanied by spikes in the pressure drop and shear stress, as the narrowing of the vessel increases its resistance to flow. Similarly, increases in cell number result in a drop in resistance and the pressure drop, which decreases the amount of shear stress present in the vessel.

## Discussion

Branched vascular networks are a heavily conserved feature across animal physiology, and this geometric configuration is vital for successful transportation of blood. The fractal-like nature of embedded vascular networks provides effective transport of nutrients and waste across the tissue domain by maximising surface area available for transport, minimising diffusion distance, and reducing the energetic cost of forcing blood flow [22–25]. Therefore, during the development of these networks an optimal level of branching must be achieved. Sprouting angiogenesis produces an initially hyperbranched network with numerous redundant flow paths, and the remodelling phase works to optimise this network by removing some branches whilst maintaining others at key locations. As of now, the exact mechanisms as to how ECs within developing networks collectively decide on what the optimal number of branches remains unknown. Franco et al. demonstrated how ECs responding to shear stress differences at vessel bifurcations is a mechanism triggering regression and loss in branching [5]. However, simulations of blood flow in developed vascular networks reveal that numerous bifurcations with shear stress differences can exist in late-stage/post-vascular remodelling [21], indicating that shear stress differences are required but not sufficient for regression and bifurcation loss. This brings us to the fundamental question of our research: what determines if a bifurcation is to be removed or preserved during flow-mediated vascular remodelling?

For a vessel to persist during remodelling, the net flux of cells entering and leaving cannot be negative. Therefore, upon reaching a vessel bifurcation, cells must finely tune migration along both possible paths if the bifurcation is to remain. Flow-migration coupling demonstrates that cells have a tendency to migrate from low to high-shear vessels [5,6,8]. However, these cells must migrate as a collective with adherens junctions intact in order to maintain fluid barrier function and prevent leakage. Therefore, the directional cues due to force transmission at adherens junctions must also play a role in influencing cell migration decisions [11,26]. However, how these migratory cues are integrated at the cellular level to promote or avoid vessel loss is unknown and currently challenging to directly prove experimentally. Our findings demonstrate that shear stress and junction force signalling interact competitively as mechanisms for determining cell migration paths at bifurcations. Furthermore, we predict that cells must integrate both cues nearly equally to maximise bifurcation stability. Dominance by one over the other leads to regression and loss of the bifurcation. The competitive nature of these two cues arises hydrodynamically from the inverse relationship between vessel diameter (and hence cell number) and resistance to flow. These findings also imply a previously unreported emergent mechanism protecting highly dynamical developmental networks against bifurcation loss: a branch that is narrowing due to a net efflux of cells will have a low junction force transmission at the bifurcation, but the decreased diameter will cause a spike in resistance and pressure drop within the vessel. This pressure spike will increase the shear stress within the vessel, attracting new cells in order to restore the vessel and prevent collapse and regression.

Our analysis of the A branch model is reminiscent of the shunt problem proposed by Pries et al. in which they describe vasculature as a coexistence of long flow pathways and short arteriovenous (AV) connections (corresponding to the distal and proximal paths in the A branch model, respectively) [27]. Pries et al. proposed the problem of these networks forming functional shunts during remodelling: assuming similar vessel diameters, the higher shear stress within the shorter AV connections will enlarge and reinforce these connections while diverting flow from the longer distal pathways. Although Pries et al. did not focus on flow-mediated migration during the remodelling process, the very same shunt problem exists within the developing vascular plexus. Indeed, we found similar functional shunt formation in our model when we set shear stress the sole factor in EC decisions at bifurcations (*α* = 1.0) and found that ECs reinforced the shorter proximal path at the expense of the longer distal path in 73% of cases. In their original analysis of the shunt problem, Pries et al. suggested that there must be an additional signal transferring information to ECs against the direction of blood flow from the distal pathways that prevents shunt formation. The authors proposed cell-cell signalling via gap junctions as a possible mechanism, but no unequivocal demonstration of this mechanism has been proposed to date. Our findings on the competitive interplay between shear sensing and junction force transmission during remodelling suggest that this additional signal that allows the distal pathways to remain intact could be force transmission through the collective endothelium during migration.

In the current study, we did not explicitly consider the molecular regulation of flow-migration coupling during angiogenic remodelling. However, several important signalling regulators of bifurcation stability have been implicated. For example, BMP-ALK1-SMAD signalling facilitates cell migration and vessel stability in low shear stress environments to prevent excessive regression [7,28]. VEGF3 modulates EC flow-migration coupling by influencing shear stress sensitivity [29]. Notch signalling facilitates polarisation against flow and artery-vein specification [30–32]. Lastly, signalling from vascular mural cells promotes vascular stability and survival [33], and the specific role of mural signalling during angiogenic remodelling and vessel regression is currently unknown. An advantage of our modelling approach is that we implicitly account for both flow- and collective-directed migratory cues and utilise a single parameter, *α*, to control the relative weight ECs place on each when selecting migration paths at bifurcations. Future work will investigate functional formulations of *α* that can capture this molecular regulation.

In conclusion, we have designed the first agent-based model of EC flow-migration coupling during developmental angiogenic remodelling in order to interrogate the cellular dynamics at bifurcations leading to vessel preservation or regression. We found that bifurcation stability can be achieved through a combination of flow-based and collective-based migratory behaviour, and these two factors can interact cooperatively or competitively depending on the scenario at the bifurcation. In cases of competition where EC migratory paths split, weighting both factors equally resulted in maximum stability and minimum loss in branching. Our findings were robust across numerous changes in the model, suggesting we have uncovered an inherent property of the physical system rather than an artificial construct of the model and its assumptions. Furthermore, our work provides a theoretical basis for the experimental investigation of bifurcation stability during angiogenic remodelling. Future theoretical work will investigate emerging behaviour in complex network environments (e.g., full vascular plexus models) and mathematical modelling of the molecular regulation behind flow-migration coupling. In particular, we are interested in instances in which the normal flow-mediated remodelling process is disrupted, leading to pathological structural abnormalities and functional shunting of vascular beds (e.g., arterio-venous malformations).

## Methods

### The flow boundary-value problem

The ABM of flow-migration coupling consists of a network of blood vessels composed of migratory ECs represented by agents. This vessel network consists of a feeding vessel serving as a flow inlet which branches into a shorter proximal branch and longer distal branch. The two branches then reconnect with each other at the draining vessel, which serves as the flow outlet. The distal branch is twice the length of the proximal branch, which is twice the length of the feeding and draining vessels (Fig 1 A). Flow, *q*, is driven by differences in pressure, *p*, along the network and pressure boundary conditions prescribed at the inlet (*p_in_*) and outlet (*p_out_*). Pressure boundary conditions were chosen to match predictions of shear stress within the capillary plexus of a developing mouse retina ranging from 1-5 Pa prior to remodelling [21]. The model simulates discrete steps in time, during which the agents move along the network (against flow). After each migration step the configuration of the network changes, so flow is recalculated and the next time step takes place in this updated flow environment, creating the coupling between migration and flow.

The vascular network was represented as a collection of *N_seg_* connected line segments with *N_node_* nodes at the segment junctions and network boundary. Pressure is assigned at each of the nodes, while flow is assigned to each segment. The length of each segment correlates to the length a migrating EC can move over the chosen length of time (*l_seg_*) based on a migration speed, *v*. The number of cells composing a vessel segment, *n*, was set to the initial value of *n*_0_ at time *t* = 0. Although the model operates solely in 2D, we approximate the diameter of the vessel lumen, *d*, in 3D as the circumference formed by wrapping the *n* cells that make up the vessel into a circle,

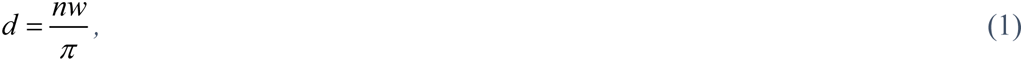

where *w* is the lateral width of an EC. Our model currently represents cells as rigid bodies with constant axial length, lateral width, and surface area. With the lumen diameter, we can then calculate the vessel conductance to flow (i.e., the amount of flow generated by a given pressure difference) as,

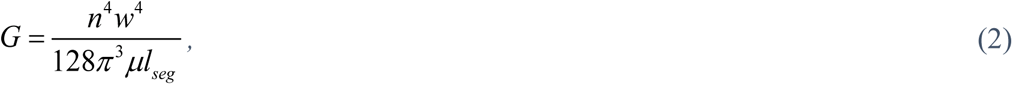

where *μ* approximates the dynamic viscosity of blood. For completeness, the vessel resistance to flow (i.e., the amount of pressure difference required to generate a given flow) is the inverse of the conductance,

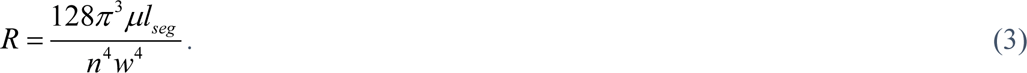

We use a flow balance equation at each node to assemble to global system of equations for the network, which we can solve for the unknown nodal pressures and therefore flow based on vessel conductivity,

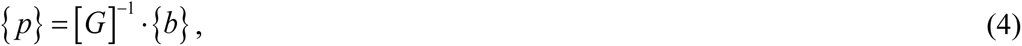

where {*p*} is the array of unknown nodal pressures, [*G*] is global conductivity matrix consisting of the individual segments’ conductivity, and {*b*} is the solution array which contains information on the pressure boundary conditions and enforces the flow balance. Vessel segments with a cell number and therefore lumen diameter of zero had their conductance set to an infinitesimally small value (1e-25 m^3^/s/Pa) as to keep [*G*] invertible. We solved the system of equations at each time step using the *numpy.linalg.solve* function, part of the *SciPy* Python library. Once we have obtained the unknown pressures, we can calculate flow through each segment as

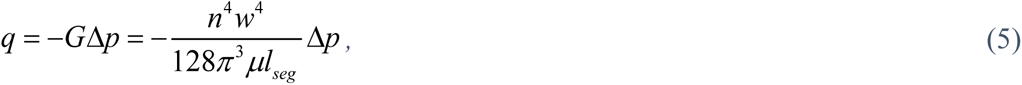

Where Δ*p* is the pressure difference between the segment’s downstream and upstream nodes. Similarly, we can calculate the wall shear stress experienced by ECs along the inner surface of the vessel lumen as,

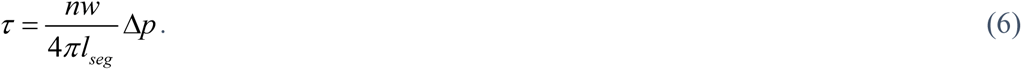

The initial state of flow and shear stress in the network prior to any migration can be found in Fig 1 B.

### EC migration, boundary conditions, and bifurcation rules

The ABM then takes a migration step within the current flow environment. For the majority of segments within the network, this simply involves moving the cells it currently has to its upstream neighbour while receiving the cells from its downstream neighbour. We included a diffusion-like intercalation component in our migration step that smooths sharp fluctuations in vessel diameter. During each migration step, if the number of cells leaving a segment is greater than the number of incoming cells (and the number of incoming cells is greater than zero), then one cell was randomly chosen to stay behind and not migrate during that step. Note that this intercalation was not implemented at vessel bifurcations to avoid interference during our analysis of EC decision behaviour at these locations.

There are several exceptions to the migration step that need to be handled specifically. The first exception is to handle cell boundary conditions, namely the case of cells migrating out at the inlet of the network as well as the condition of cells entering the network at the outlet. We explored two different boundary conditions for cell migration: a periodic condition, and a Dirichlet condition. In the periodic condition, any cells migrating out at the inlet re-enter the network at the outlet, keeping the total number of cells within the network constant throughout the simulation. In the Dirichlet condition, the number of incoming cells at the outlet was kept fixed at the initial cell number *n*_0_, which means only so many cells exciting at the inlet were allowed to enter the outlet at any time. This causes the total number of cells within the simulation to fluctuate over time in order to match this condition. This manuscript will cover the periodic conditions in the main text, for information on results using the Dirichlet conditions please see the Supporting Material (S9 Fig).

The remaining migration exceptions involve the two bifurcations within the network. The bifurcation at the left of the network (near the inlet) is a flow-divergent bifurcation which means EC migration paths converge. At this bifurcation, incoming cells from both branches only have one choice to migrate against the flow which is to join at the feeding vessel before exciting through the inlet. The bifurcation on the right of the network (near the outlet) presents a much more interesting case. This is a flow-convergent bifurcation where EC migration paths diverge: cells approaching this bifurcation have two choices available when migrating against the flow. Due to the difference in path lengths between the proximal and distal branch, the flow and shear stress in the proximal segment (hence referred to as branch 1) is higher than the distal segment (hence referred to as branch 2).

Since the actual mechanisms regulating EC migration at vessel bifurcations during flow-mediated remodelling are unknown, we designed and implemented various “bifurcation rules” for our agents upon approaching the flow-convergent bifurcation and observed the emergent outcomes on the remodelled vessel network. In our first bifurcation rule (BR 1), shear stress is the only determinant of which branch the ECs choose, such that

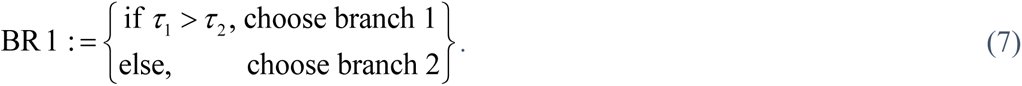

In the second bifurcation rule (BR 2), ECs choose the branch that forms the shallowest angle with the parent branch such that cells choose the branch that requires the least change in direction,

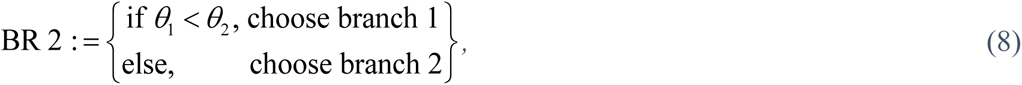

where *θ*_1_ and *θ*_2_ are the angles between the parent segment and branches 1 and 2, respectively.

The next set of rules utilise a stochastic description of EC behaviour at bifurcations, in which each cell at the bifurcation generates a random number, *r*, between 0 and 1 and chooses a branch based on this number compared to the probability of entering branch 1 (*P*_1_) or branch 2 (*P*_2_), where *P*_2_ = 1− *P*_1_. In the third bifurcation rule (BR 3), we assigned both branches equal probability *P*_1_ = *P*_2_ = 0.5, which remains constant throughout the simulation,

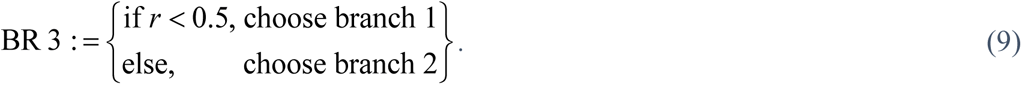

In the fourth bifurcation rule (BR 4), we use a similar mechanism but with unequal probabilities *P*_1_ = 0.7, *P*_2_ = 0.3,

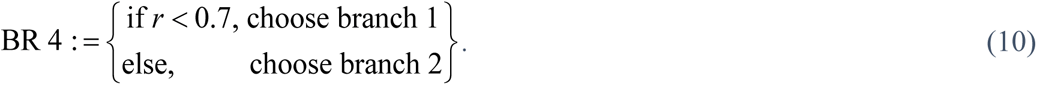

In the final bifurcation rule (BR 5), we sought to update the branch probability over time to dynamically couple it with EC migration. In this rule we utilise a similar scholastic mechanism,

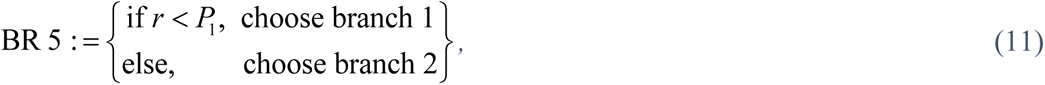

where each branch probability is constantly updated via weighted average of the probability due to shear stress (*P_τi_*) and the probability due to cell number (*P_ni_*),

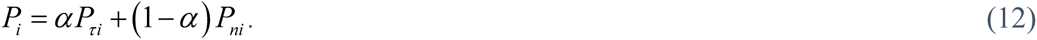

The shear stress and cell number probabilities come from the ratio of shear stress and cell number between the two branches, respectively,

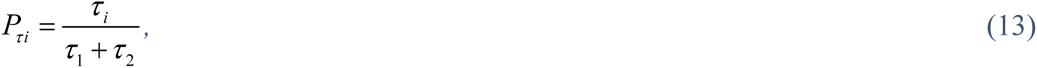

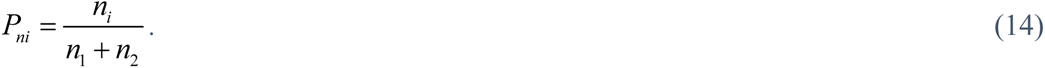

Note that Eq 6 can be substituted into Eq 13 to express the shear stress probability as the ratio of the multiplicative combination of cell number and the pressure difference over the branch segment,

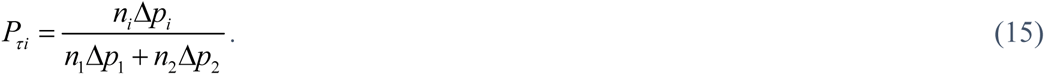

### Overview of simulations and data analysis

For a full list of all parameters and variables with the model as well as values used in the simulations, please see Table 1. We performed several simulations for each bifurcation rule and quantified the results in order to characterise the impact each rule had on network remodelling. Mean diameter of the proximal and distal branch was measured by taking the mean of the diameter across all adjacent segments in that branch. The same number of segments was included in this mean for both the proximal branch and distal branch, even though the distal branch was twice in length. The first two bifurcation rules, BR 1 and BR 2, do not depend on any randomness so only a single simulation was performed. For the remaining rules, we varied the seed number for the random number generator accordingly. We generated a list of 1000 seed numbers between 1 and 10^9^ using the *randint* function in the *random.py* Python library. Each simulation utilised a different number from this list as its seed number, and the same list was used across all other parameter and rule changes.

**Table 1.** Glossary of constants and variables with the model, values and units.

| constants |  |  | variables |  |
| --- | --- | --- | --- | --- |
| name | description | value & units | name | description (units) |
| $N_{seg}$ | number of segments | 40 | $p$ | blood pressure (Pa) |
| $N_{node}$ | number of nodes | 41 | $\Delta p$ | pressure difference (Pa) |
| $l_{seg}$ | length of segment | 10 $\mu\text{m}$ | $q$ | blood flow ($\mu\text{L/day}$) |
| $v$ | migration speed | 3 $\mu\text{m/hr}$ | $\tau$ | wall shear stress (Pa) |
| $w$ | lateral width of EC | 5 $\mu\text{m}$ | $n$ | cell number |
| $\Delta t$ | time step size | 3 $\frac{1}{3}$ hr | $\alpha$ | branching mechanism component weight |
| $p_{in}$ | inlet pressure | 100 Pa | $\theta$ | branching angle |
| $p_{out}$ | outlet pressure | 0 Pa | | |
| $\mu$ | dynamic viscosity of blood | 0.0035 Pa-s | | |

In the basic stochastic rules BR 3 and BR 4, we only ran simulations using the first 10 numbers as this was sufficient to characterise trends outside of randomness in those cases. When assessing the parameter *α* in the mechanistic bifurcation rule BR 5, we varied *α* across its range from 0.0 to 1.0 at intervals of 0.01 and ran the full list of 1000 seed numbers for every value of *α*. We monitored the cell number in each of the segments composing the flow-convergent bifurcation and when the cell number in either of those segments dropped to zero, we considered the bifurcation lost. We collected the number of simulations with bifurcation loss over time and normalised by the number of simulations (*M* = 1000) in order to obtain the percent bifurcation loss for that value of *α*. When analysing the branch probability, we took the shear stress and cell number from both of those branches to calculate the probability as a function of time.

Finally, we performed additional simulations not included in the main body of the text in order to ensure that our findings were not specific to any particular configuration or set of conditions. This included prescribed inlet flow boundary conditions, Dirichlet cell boundary conditions, distal paths with increased resistance in the A branch model, and a model of a single flow-convergent bifurcations (Y branch model). For information on the results of these additional simulations, please see the Supporting Material.

## Acknowledgements

We would like to graciously acknowledge our funding as part of a Foundation Leducq Transatlantic Network of Excellence (17 CVD 03, https://www.mdc-berlin.de/leducq-attract).

## Supporting Information

For all videos in the Supporting Material, we recommend VLC media player (https://www.videolan.org/vlc/index.en-GB.html)

**S1 File. *S1_ABM_A_branch_BR_1.mp4*.** Simulation in the A branch model using Bifurcation Rule 1, in which cells choose the branch with largest shear stress.

**S2 File. *S2_ABM_A_branch_BR_2.mp4*.** Simulation in the A branch model using Bifurcation Rule 2, in which cells choose the branch that requires the least change of direction.

**S3 File. *S3_ABM_A_branch_BR_3.mp4*.** Simulation in the A branch model using Bifurcation Rule 3, in which cells randomly choose a branch with equal preference.

**S4 File. *S4_ABM_A_branch_BR_4.mp4*.** Simulation in the A branch model using Bifurcation Rule 4, in which cells randomly choose a branch with unequal preference (favouring the high-flow proximal branch).

**S5 File. *S5_ABM_A_branch_BR_5_alpha_0.00.mp4*.** Simulation in the A branch model using the Mechanistic Bifurcation Rule (BR 5) with *α* set to 0.0; cells consider only cell number when choosing a branch.

**S6 File. *S6_ABM_A_branch_BR_5_alpha_1.00.mp4*.** Simulation in the A branch model using the Mechanistic Bifurcation Rule (BR 5) with *α* set to 1.0; cells consider only shear stress when choosing a branch.

**S7 File. *S7_ABM_A_branch_BR_5_alpha_0.45.mp4*.** Simulation in the A branch model using the Mechanistic Bifurcation Rule (BR 5) with *α* set to 0.45; cells consider both cell number and shear stress differences when choosing a branch.

**S8 Fig.**
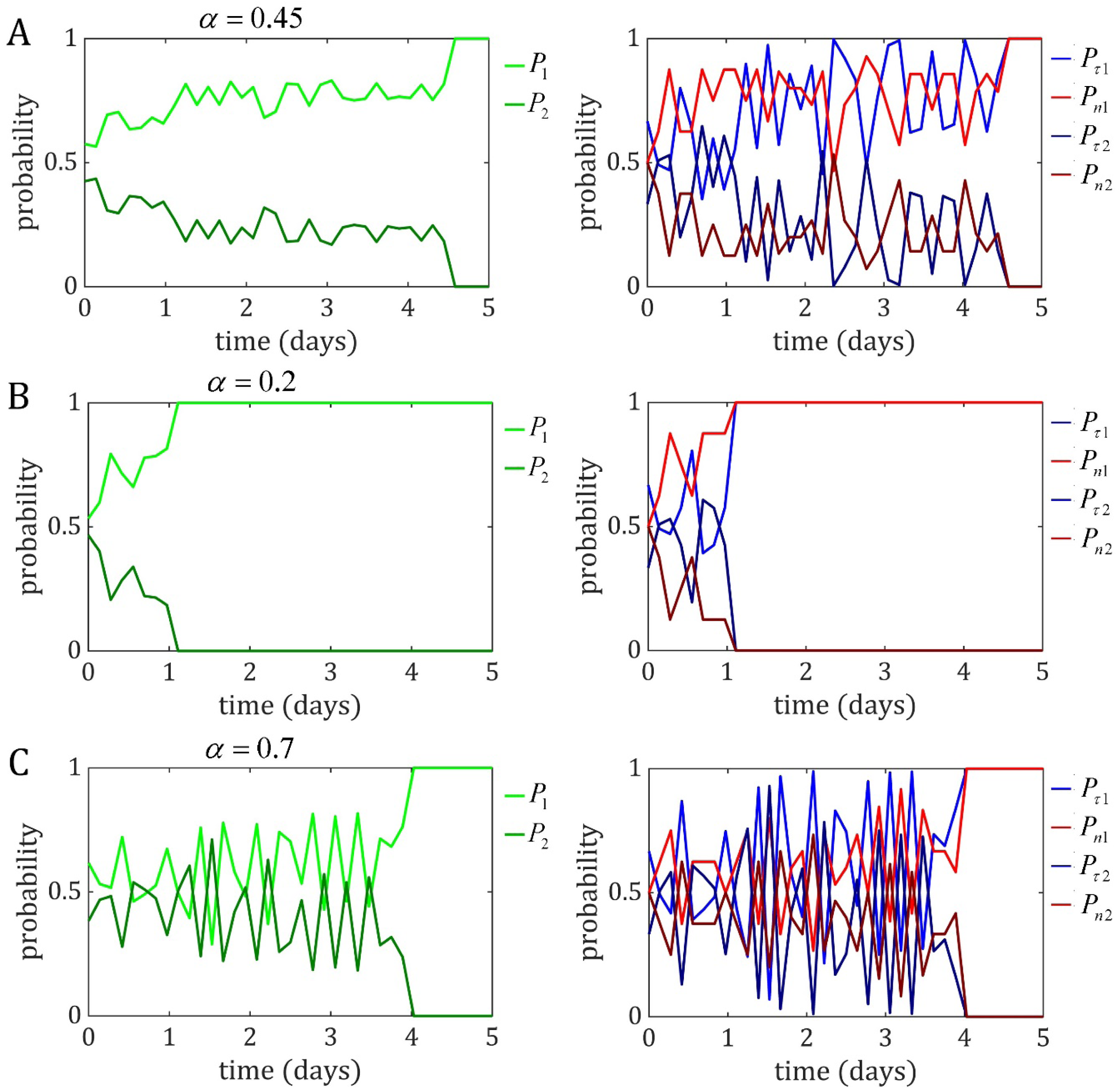
Examples of probabilities from simulations that exhibited bifurcation loss. Each result was obtained from the same random seed number with *α* values of 0.45 (A), 0.2 (B), and 0.7 (C). In all cases, there were temporary oscillations between shear stress and cell number probability while the bifurcation remained stable, but in each case a branch probability reached maximum likelihood (i.e., equal to 1) and the bifurcation was lost.

**S9 Text. Stability is robust but does vary across changes in boundary condition, network geometry, and shear stress difference**

**Results**

The final task of our study was to demonstrate that the stability behaviour described previously is not exclusive to our pressure-driven A branch model with periodic cell conditions. Hence, we performed similar stability analysis across different boundary conditions and vessel network geometries. The first variation of the A branch model included a prescribed flow condition at the inlet as opposed to prescribed pressure (S10 File). In this model, the pressure at the inlet varies in order to set a prescribed flow at the inlet, which was set to the same initial incoming flow in the pressure-driven model. Prescribing inlet flow instead of pressure had no effect on the stability at the flow-convergent bifurcation, as bifurcation loss vs. *α* over time was identical to the pressure-driven results (S11 Fig). Next, we sought to determine the impact on the EC boundary conditions on stability by holding the number of incoming cells constant, meaning the total number of ECs within the network varied in order to match this condition (as opposed to the periodic condition where the total number of ECs was held constant) (S12 File). Simulations with this condition resulted in a similar asymmetric saddle-shape while sweeping over the range of *α*; however, the global minimum of bifurcation loss was shallower (i.e., less stable) when compared to the periodic cell conditions (S13 Fig). In our final variation of the A branch model, we sought to determine how the initial shear stress difference at the bifurcations impacted stability. In the original A branch model, the shear stress difference at the bifurcation had an initial ratio of 2:1, proximal to distal, due to the distal path being twice the length (and hence twice the resistance) of the proximal path. We therefore created a new version of the A branch model in which the distal path was 10× the length of the proximal path, creating an initial shear stress ratio of 10:1. Stability analysis with this model resulted in a similar asymmetric saddle to the 2:1 case, although the global minimum was shallower (less stable) and shifted slightly to the left centred around *α* = 0.375 (S14 Fig).

The A branch model is a basic representation but still includes some “network effects” as flow is split at the flow-divergent bifurcation before re-joining at the flow-convergent bifurcation. This model has some advantages, specifically in that cell behaviour at the flow-convergent bifurcation doesn’t propagate to the flow inlet as cells in both paths combine at the flow-divergent bifurcation. However, varying the shear stress difference at the bifurcation in this model is not straightforward and requires a change in geometry (i.e., change in path lengths between the two branches). Thus, we created a simplified model of a flow-convergent bifurcation (hence referred to as the Y branch model) in which incoming flow from a left and right branch combine at a single bifurcation before exiting via the outlet (S15 File). Using this model, we can set the inlet pressure in both the left and right branches to be equal in order to achieve a 1:1 shear stress ratio (i.e., no shear stress difference) at the bifurcation (S16 Fig). Stability across the range of a in this model was very similar to the pressure-driven A branch model, with a global minimum across values between 0.3 and 0.6. The branch probabilities in this model initialise at 0.5 as the cell number and shear stress in both branches is the same and exhibit similar competitive oscillations which result in bifurcation stability (S17 Fig). Finally, we implemented the Y branch model with inlet flow conditions in both branches that allows us to vary the initial shear stress difference at the bifurcation at ratios of 1:1, 1:10, and 1:20 (S18 File, S19 File, S20 File). Each of these cases resulted in a similar saddle shape in the stability surface, although this saddle became shallower (less stable) and shifted to the left (towards *α* = 0.0) as the shear stress difference increased (S21 Fig). While there was no difference in mean diameter between the two branches with a shear ratio of 1:1, increasing the ratio between shear stress led to larger differences in diameter between the two branches (S22 Fig).

**Discussion**

Our stability findings were robust across changes in boundary condition, vessel geometry, and shear stress initialisation at the bifurcation, strongly suggesting that our findings are truly inherent to physical system we are representing and not a manufactured outcome of any one particular model configuration. Inlet flow conditions made no impact of stability at the flow-convergent bifurcation in the A branch model, most likely due to the face that cells from both branches recombine at the flow-divergent bifurcation rendering the inlet free from EC behaviour at the flow-convergent bifurcation. In the inlet flow version of the Y branch model, which does not include this recombining effect, we found slightly different results but the classic asymmetric saddle shape of stability remained intact. Changing the initial conditions for shear stress (and hence shear stress probability) could affect the nature of the competitive oscillations between the two components; however, when we varied these initial conditions (either by changing the resistance of the distal branch in the A branch model or the flow ratio between branches in the Y branch model) the classic asymmetric saddle shape of stability was maintained albeit more shallow and shifted to the left towards *α* = 0.0. With an increased initial shear stress difference, simulations that favoured junction forces resulted in fewer bifurcations lost, although results were still not as stable when compared to simulations with less of a shear stress imbalance. Changes in the boundary condition governing the number of incoming cells had the greatest impact on bifurcation stability. Simulations which held the amount of incoming cells constant exhibited a general decrease in the total number of cells within the system, and although we found the same asymmetric saddle shape the bifurcation was inherently more unstable with 40% of simulations losing the bifurcation by day 5 of migration (as compared to around 20% found with the Periodic cell condition). These findings suggest that one of the most significant regulators of bifurcation preservation may be the level of new incoming cells: a steady source of new cells greatly enhances the chance of a bifurcation remaining during remodelling. Likewise, a bifurcation which is experiencing reduced or disrupted levels of incoming cells may be more prone to instability and regression. At this point it is unclear what acts as the source of cells in the developing network in vivo. Proliferation may be isolated to the sprouting front, meaning new ECs must arrive via travelling down the network to the remodelling zone. Additionally, large vessels such as veins could also act as a reservoir of new cells. More experimental evidence is required in characterising the source of new ECs in the developing retina in order to investigate the matter further.

**S10 File. *S10_ABM_A_branch_BR_5_alpha_0.45_inlet_flow.mp4*.** Simulation in the A branch model (BR 5, *α* = 0.45) with the inlet flow boundary condition.

**S11 Fig.**
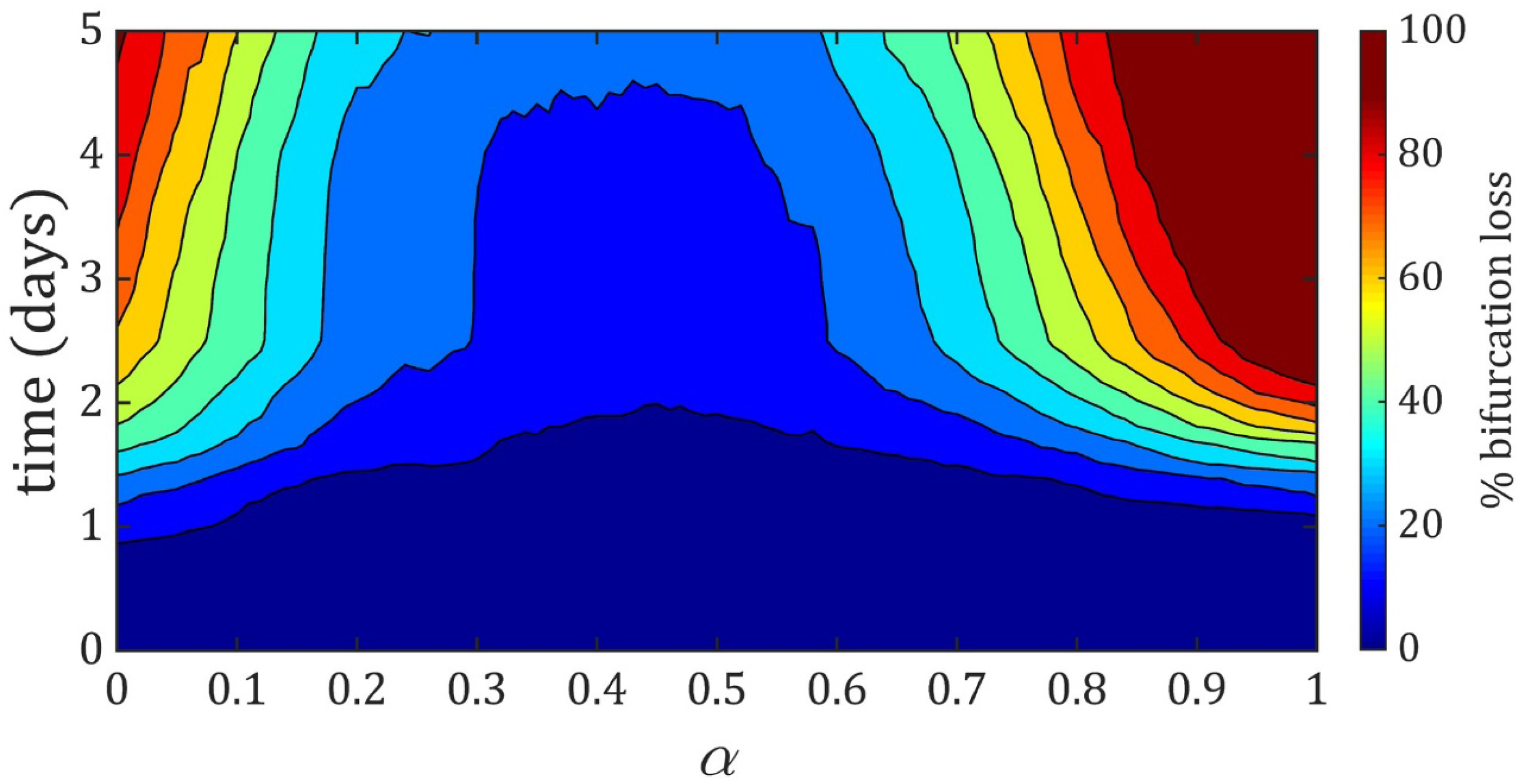
Bifurcation stability in the A branch model with inlet flow condition. Similar stability analysis as found in Fig 3 with the A branch under inlet flow boundary conditions. Inlet flow was prescribed to match the same initial incoming flow in the pressure-driven formulation. Stability results were identical in the inlet flow version of the model when compared to the pressure-driven formulation.

**S12 File. *S12_ABM_A_branch_BR_5_alpha_0.45_Dirichlet_cell_BCs.mp4*.** Simulation in the A branch model (BR 5, *α* = 0.45) with pressure-driven flow and Dirichlet cell boundary conditions, in which the number of cells entering the domain was held constant.

**S13 Fig.**
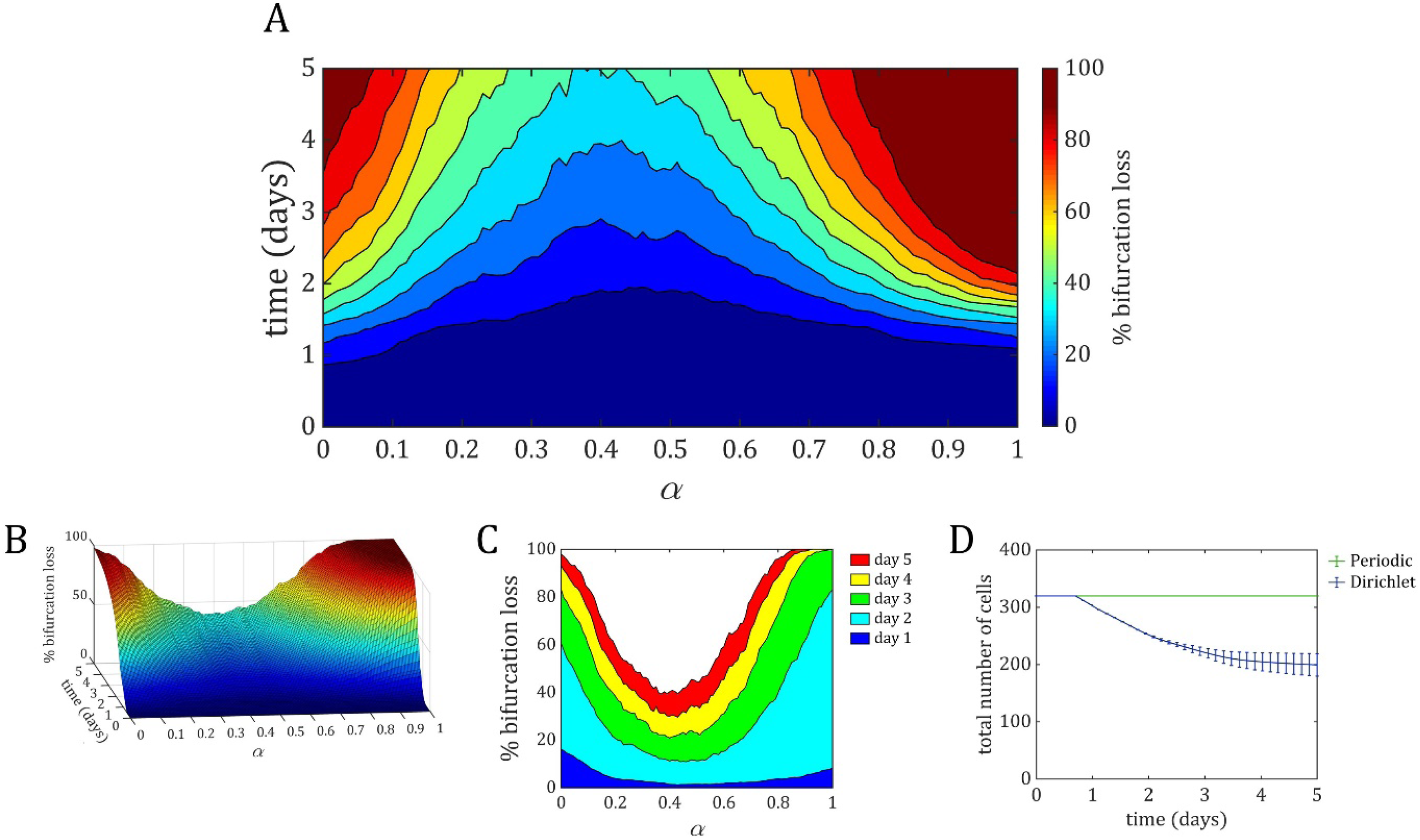
Bifurcation stability in the A branch model with Dirichlet cell boundary conditions. A similar sweep through the values of *α* to Fig 3 but with Dirichlet cell boundary conditions applied to the model rather than periodic boundary conditions. These boundary conditions enforce that in number of cells incoming to the network (at the flow outlet) was held constant throughout the simulation. This results in the total number of cells within the domain changing over time as this condition is enforced. Simulations using the Dirichlet boundary condition resembled those of the periodic boundary condition but were generally less stable. (A) The contour plot of bifurcation stability vs. *α* over time shows a similar global minimum of stability around *α* = 0.4, but even within this stable region more simulations lost the bifurcation at earlier points in time compared to simulations with the periodic boundary condition. (B) The surface formed by bifurcation stability vs. *α* over time resembled a similar asymmetric saddle shape but was much steeper, especially in the more stable region, indicating that this region was more unstable when compared to the periodic boundary conditions. (C) Similar to the periodic boundary condition case, the majority of bifurcations were lost during day 2 of migration. However, simulations with the Dirichlet condition lost more bifurcations at later days (days 3, 4, 5) when compared to periodic simulations. (D) The total amount of cells in the domain for 1000 simulations at *α* = 0.45 for both the Periodic and Dirichlet cell condition. The total number of cells remains constant with the Periodic condition while decreasing over time with the Dirichlet condition.

**S14 Fig.**
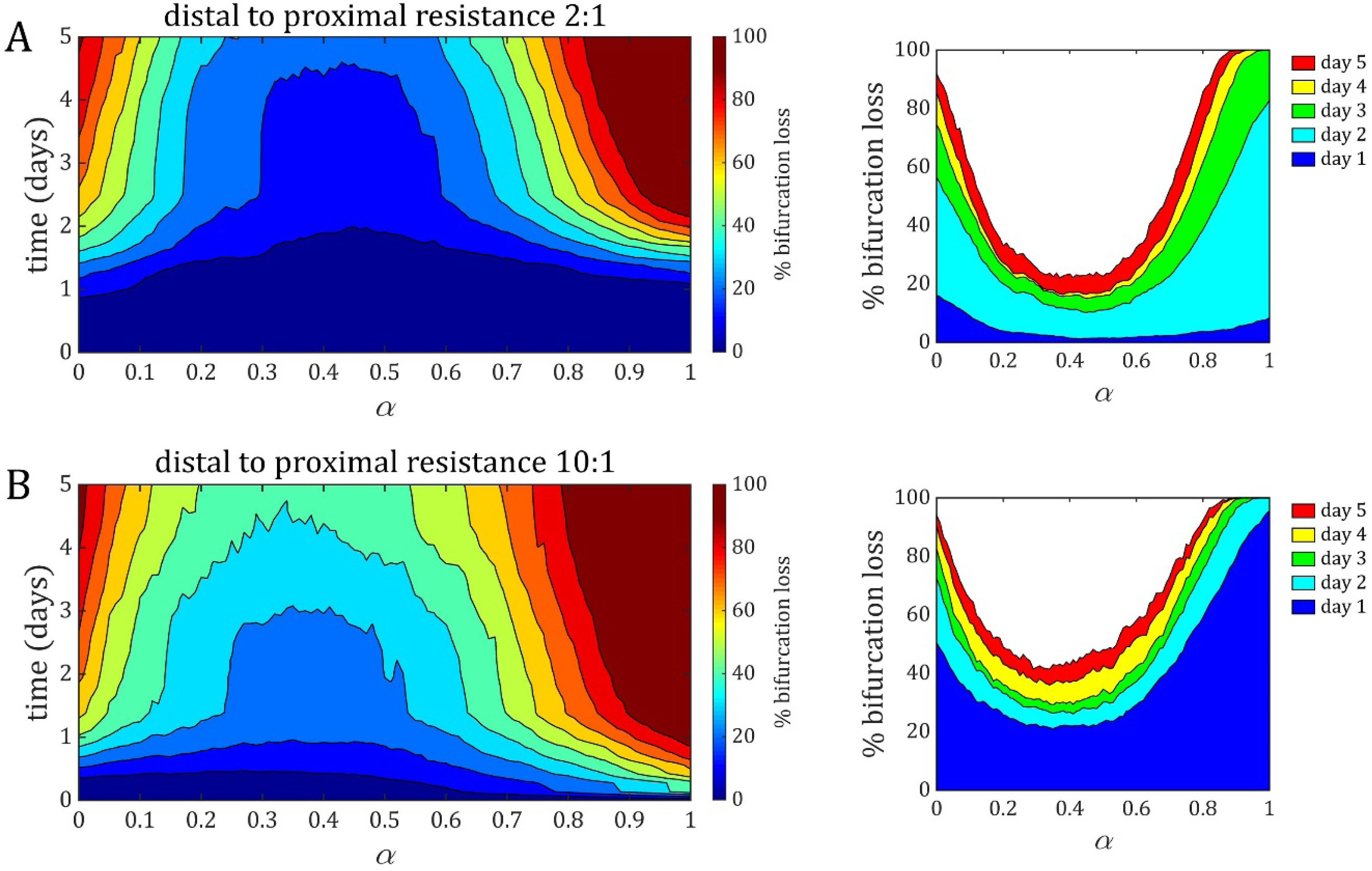
Varying the initial shear stress difference at the bifurcation in the A branch model by increasing the length of the distal branch. (A) Stability results from the original formulation of the A branch model which includes an initial 2:1 shear stress difference at the flow-convergent bifurcation. (B) Stability results from a version in the model in which the distal branch was 10× longer than the proximal path, resulting in a 10:1 shear stress difference at the bifurcation. We found a similar saddle shape in stability vs. *α* over time, although much shallower when compared to the original results indicating that bifurcation occurred more readily with the larger shear stress difference.

**S15 File. *S15_ABM_Y_branch_BR_5_alpha_0.45.mp4.*** Simulation in the Y branch model (BR 5, *α* = 0.45) with initially equal flow in both branches (and hence no initial difference in shear stress at the bifurcation).

**S16 Fig.**
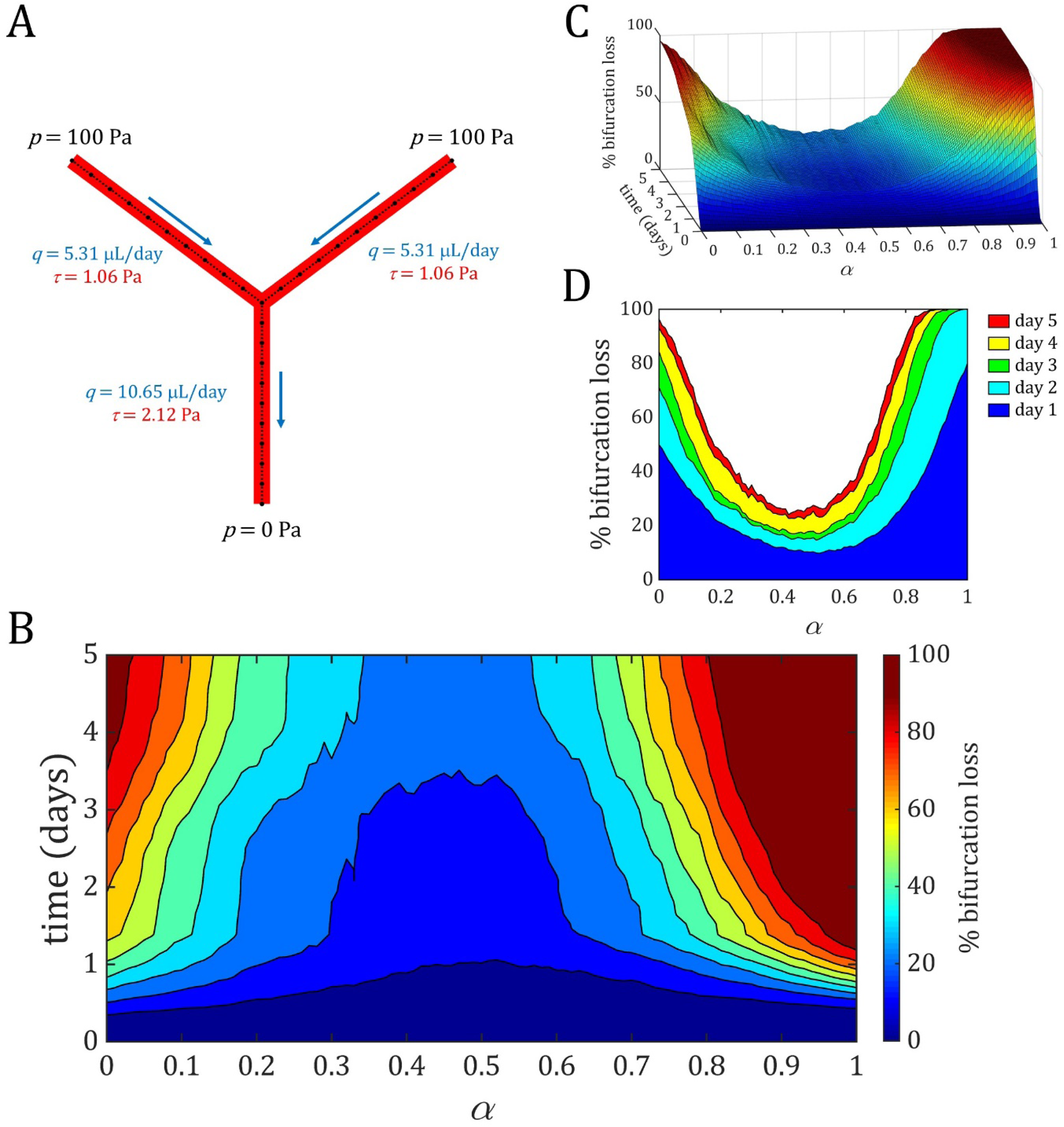
Stability in the Y branch model with no initial shear stress difference at the bifurcation. (A) Schematic of the Y branch model with pressure boundary conditions set equal at both inlets, resulting in a 1:1 shear stress at the bifurcation. (B-D) Stability vs. *α* over time resulted in a similar saddle shape with global minimum centred around *α* = 0.45 when compared to the A branch model.

**S17 Fig.**
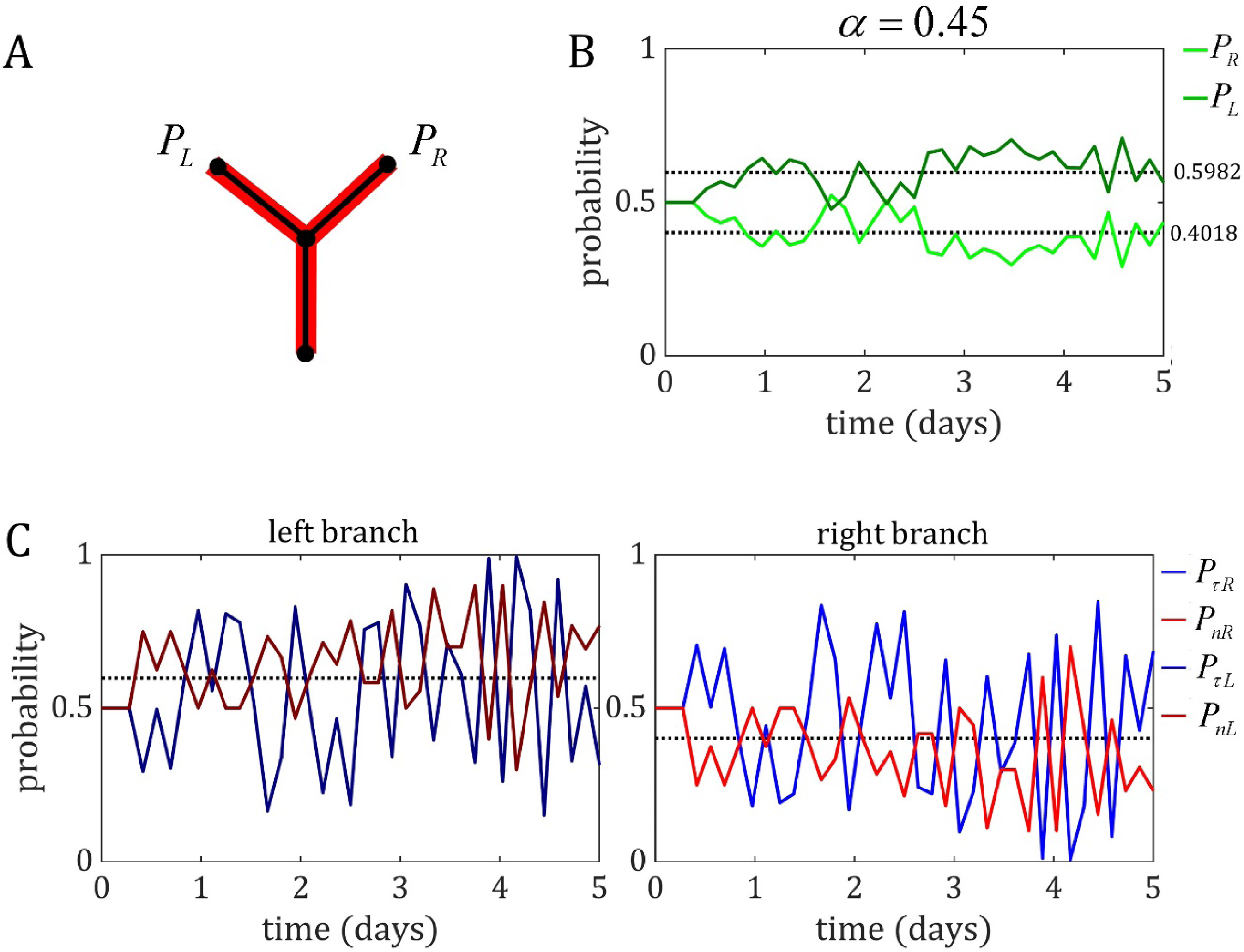
Similar competitive oscillations between shear stress and junction force probability found in the Y branch model without initial shear difference. (A) Probability of cells choosing the left branch (*P_L_*) or right branch (*P_R_*) upon encountering the bifurcation during migration. (B) Probability of each branch initialises at 0.5 (as both branches have the same initial amount of WSS). During the simulation, the branch probabilities tended to separate slightly, favouring one branch over the other. (C) Similar competition between shear stress and cell number probability produces stability at the bifurcation.

**S18 File. *S17 _ABM_Y_branch_BR_5_alpha_0.45_flow_ratio_1to1.mp4*.** Simulation in the Y branch model (BR 5, *α* = 0.45) with inlet flow conditions and a 1:1 flow ratio between the left and right branch.

**S19 File. *S18 _ABM_Y_branch_BR_5_alpha_0.45_flow_ratio_1to10.mp4*.** Simulation in the Y branch model (BR 5, *α* = 0.45) with inlet flow conditions and a 1:10 flow ratio between the left and right branch.

**S20 File. *S19 _ABM_Y_branch_BR_5_alpha_0.45_flow_ratio_1to20.mp4*.** Simulation in the Y branch model (BR 5, *α* = 0.45) with inlet flow conditions and a 1:20 flow ratio between the left and right branch.

**S21 Fig.**
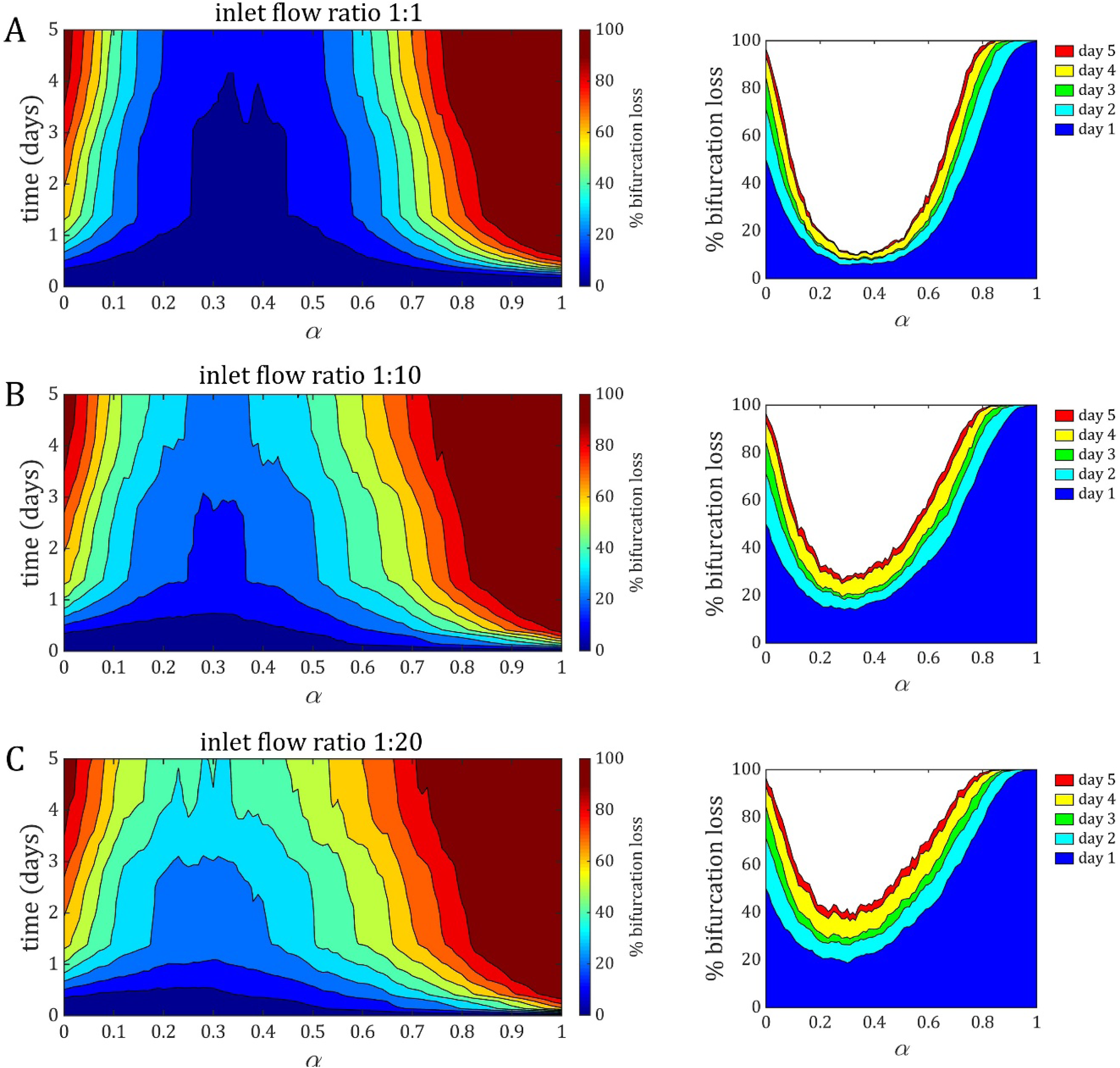
Stability in the Y branch model with inlet flow conditions whilst varying the initial shear stress ratio at the bifurcation. Inlet flow conditions in the Y branch model were used to create initial shear stress ratios between the left and right branch of (A) 1:1, (B) 1:10, (C) 1:20. In general, that stability saddle shifted upward and to the left (towards *α* = 0.0) as the initial shear stress difference at the bifurcation increased.

**S22 Fig.**
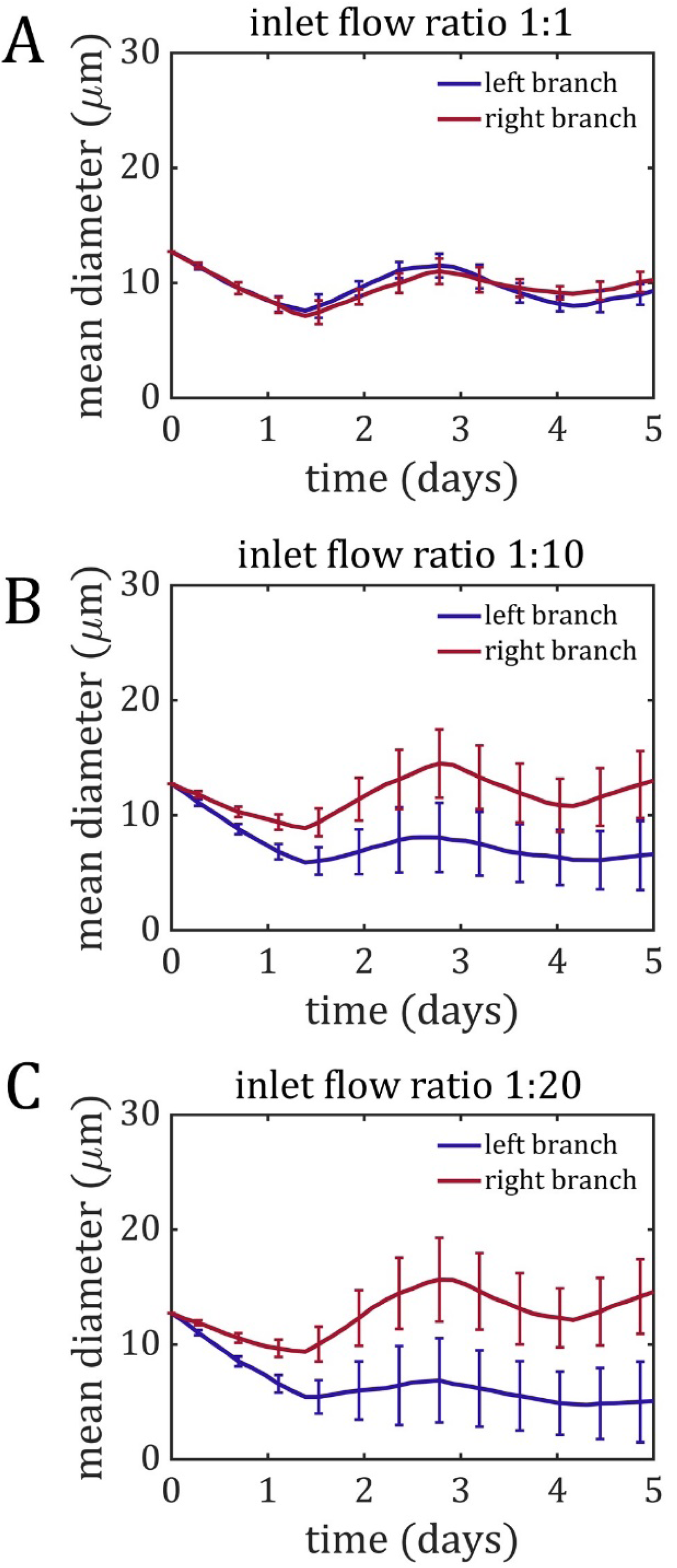
Difference in stabilised diameters between the two branches increases with inlet flow ratio. Mean diameter of the left branch (blue) and right branch (red) in the Y branch model with shear stress differences of (A) 1:1, (B) 1:10, and (C) 1:20. The difference in diameter between the two branches increased as this initial shear stress difference increased.

